# SALP, a new single-stranded DNA library preparation method especially useful for the high-throughput characterization of chromatin openness states

**DOI:** 10.1101/206854

**Authors:** Jian Wu, Wei Dai, Ling Wu, Jinke Wang

## Abstract

Based on a novel kind of single strand adaptor (SSA), this study developed a new method to construct next-generation sequencing (NGS) library, named as SALP, representing Single strand Adaptor Library Preparation. The key creativity of the method lies in the design and verification of a special adaptor that can be efficiently linked to the 3′ end of single-stranded DNA, which is a double-stranded oligonucleotide with a 3′ overhang of 3 random nucleotides. This method can start with the denatured DNAs or chromatins fragmented by different methods such as Tn5 tagmentation, enzyme digestion and sonication. When applied to Tn5-tagmented chromatin, SALP overcomes the key limitation of the current ATAC-seq method and develops a high-throughput NGS library construction and sequencing approach, SALP-seq, which can be used to comparatively characterize the chromatin openness state of multiple cells simply and unbiasly. In this way, the comparative chromatin openness states of four different cell lines, including GM12878, HepG2, HeLa and 293T, were successfully characterized. This study also demonstrated that SALP-seq could characterize the chromatin openness states with 10^5^ to 500 cells, indicating the high sensitivity of SALP-seq in characterizing chromatin state of cells. SALP should have wide applications in the future biological sciences and biomedicine.

## 1. Introduction

Since next-generation sequencing (NGS) appeared on the marketplace in 2005,^1^ this technology has changed the way we think about scientific approaches in basic, applied and clinical research.^2^ With the innovative methodological and computational developments, NGS platforms have facilitated an explosion on biological knowledge over past years.^3^ As the broadest application of NGS, the re-sequencing of human genomes dramatically enhance our understanding of how genetic differences affect health and disease.^2^ Compared with the Sanger sequencing, the key procedural difference in NGS methods is the construction of sequencing library.^3^ As the productivity of NGS platforms continues to grow with hardware and software optimization, the construction of sequencing libraries became the bottleneck of the using of this technology.^4^ To address the issue, various library construction methods have been increasingly explored, in which included several interesting methods starting from single-stranded DNA (ssDNA).^5–8^ However, these methods are still prevented from widely applications by their intrinsic shortcomings.

The current standard method for constructing NGS libraries starts from double-stranded DNA (dsDNA), which is entirely *in vitro* and typically includes fragmentation of DNA (sonicated or enzymatically digested), end-polishing, ligation of adaptor sequences, gel-based size-selection, and PCR amplification.^4^ With tedious procedure, many steps of the typical protocol are suboptimal, and results in considerable sample loss. To cover these shortcomings, a NGS library construction method based on DNA tagmentation was developed. In this method, two mosaic end (ME) adaptors that harbor the annealing sites of two primers were firstly complexed with hyperactive derivative Tn5 transposase to form transposome, which then tagmented DNA into tagments with adaptors at their 5′ ends.^4^ Finally, the DNA tagments were amplified with specific primers by using PCR of limited cycles, which produced the DNA library compatible with massively parallel sequencing. However, when this method was performed to create dual-tagged libraries, only part of tagments could be amplified with two different library constructing primers correctly and sequenced successfully, even suppression PCR was performed to increase the percentage of fragments with correct structure.^9^

The processes of development and differentiation of mammals are regulated by the constant changing of binding interaction between DNA-binding proteins and chromatin.^10^ The chromatin provides the restricted access of transcription factors (TFs) to their DNA binding sites in a highly cell-type-specific manner.^10–12^ As a result, numbers of researches focused on the openness of chromatin.^13–16^ For this reason, as a rapid and sensitive method to capture the open chromatin sites, the transposase-accessible chromatin using sequencing (ATAC-seq) was developed and has been widely used to research the chromatin openness state in kinds of situation.^17–20^ The mapping of chromatin openness state is beneficial for finding the key regulators driving disease. For example, the transcription factor AP1 was identifies as a transcriptional regulator in oesophageal adenocarcinoma (OAC) by profiling open chromatins in OAC-derived cell lines and patient-derived OAC material with ATAC-seq.^21^ However, ATAC-seq was limited by the intrinsic shortcoming of its tagmentation-based library construction.

Here we report a novel and versatile library construction method that can be used to capture the chromatin openness as ATAC-seq, which can construct the sequencing library with DNA samples sheared by different fragmentation methods. We named the method as SALP, which stands for the Single strand Adaptor Library Preparation. When performing the tagmentation-based library construction, dependent on the novel barcoded Tn5 adaptors (BTAs), and the method could pool the different tagmented chromatins of various cells together, which could then be taken as one DNA sample for subsequent library construction. The DNA sample was successively undergone a single strand adaptor ligation, an elongation and T adaptor ligation. The dual adaptor-ligated DNA was finally amplified with NGS compatible primers by using PCR of limited cycles. With this method, we captured high-quality chromatin openness states of GM12878, HepG2, HeLa and 293T cell lines in an unbiased and high-throughput format, which is helpful for comparatively characterizing the chromatin openness states of various cells or particular interested regulatory regions such as promoters and enhancers. With this method, we characterized the chromatin openness states of HepG2 from 10^5^ to 500 cells, indicating the method had high sensitivity in characterizing chromatin state of cells.

## 2. Material and methods

### 2.1. Cultivation of cells

HeLa, HepG2 and 293T cells were grown in Dulbecco's Modified Eagle's Medium (DMEM, GIBCO), GM12878 cells were cultured in RPMI 1640, which were all supplemented with 10% fetal bovine serum (FBS, GIBCO) with 100 μg/mL streptomycin and 100 units/mL penicillin at 37°C in 5% CO_2_.

### 2.2. Preparation of various adaptors

All oligonucleotides were synthesized by Sangong Biotech (Shanghai) (Supplementary Table 1). For preparing the barcoded Tn5 adaptors (BTAs), barcode and ME oligos were dissolved in ddH_2_O at the concentration of 20 μM, and then mixed in equimolar in PCR tube. For preparing single strand adaptors (SSAs), SSA-PN and SSA-PNrev oligos were dissolved in ddH_2_O at the concentration of 100 μM, and then mixed in equimolar in PCR tube. For preparing T adaptor, TOA and TOArev oligos were dissolved in ddH_2_O at the concentration of 100 μM, and mixed in equimolar in PCR tube. Finally, all oligo mixtures were denatured in the water bath for 5 minutes at 95 °C and gradually cooled to 25 °C for annealing into various adaptors.

### 2.3. Preparation of Tn5 transposome

Briefly, 4 μL of BTA (10 μM) was mixed with 2 μL 10× TPS, 1 μL Tn5 transposase and 13 μL H_2_O according the instruction of the Tn5 transposase (Robust Tn5 Transposase, Robustnique Corporation Ltd.). The reaction was gently mixed at 25 °C for 30 min to generate Tn5 transposome. The transposome was stored at -20 °C for later use.

### 2.4. Fragmentation of genomic DNA

The genomic DNA (gDNA) was extracted with phenol:chloroform from HepG2 cells. For tagmentation, 50 ng of gDNA was tagmented in a 30-μL reaction containing 1×LM buffer, 3 μL Dimethylformamide (DMF) and 4 μL Tn5 transposome at 55 °C for 15 min. The tagmented gDNA was purified with the MinElute PCR Purification Kit (QIAGEN, 28004). For Hind III digestion, 1 μg of DNA was digested in a 50-μL reaction containing 1× FastDigst Buffer and 5 μL FastDigest Hind III (Thermo Fisher, ER0501) at 37 °C overnight. For sonication, 1 μg of gDNA was sonicated by using the BRANSON sonicator with 70% power, 20s on and 20 s off for 20 cycles. The fragmented gDNAs were denatured at 95 °C for 5 min and chilled on ice for 5 min, which was then run with 1.5% agarose gel. The gDNA fragments of 200-1000 bp were recovered with the QIAquick Gel Extraction Kit (QIAGEN, 28704).

### 2.5. Optimization of single strand adaptor of SALP

To generate Illumina compatible sequencing libraries and improve the ligation efficiency, we designed and used single strand adaptors (SSAs) with 3′ overhang of 4 different length (1–4 random nucleotides). For SSA ligation, 12.5 ng of tagmented HepG2 gDNA was denatured at 95 °C for 5 min and chilled on ice for 5 min. The denatured gDNA was ligated with variant SSAs in a 10-μL reaction containing 1 μL of T4 DNA ligase (NEB, M0202L), 1× T4 DNA ligase buffer and 1 μL of SSA (5 μM) at 16 °C for 60 min. Then the reaction was mixed with equal volume of 2× rTaq mix (Takara) and incubated at 72 °C for 15 min. Finally, the gDNA was purified with 1.2× Ampure XP beads (Beckman Coulter) and amplified in a 50-μL PCR reaction containing 25 μL of NEBNext^®^ Q5^®^ Hot Start HiFi PCR Master Mix (NEB, M0543S), 1 μL of NEBNext Universal PCR Primer (10 μM), and 1 μL of NEBNext Index Primers (10 μM). The PCR program was as follows: (i) 98 °C for 5 min; (ii) 98 °C for 10s; 65 °C for 30s; 72 °C for 1 min; 18 cycles; (iii) 72 °C for 5 min. The PCR products was run with agarose gel and the DNA fragments of 300-1000 bp were extracted with QIAquick Gel Extraction Kit.

The prepared library was analyzed with cloning sequencing. The extracted DNA was mixed with the equal volume of 2× rTaq mix (TaKaRa, R001A) and incubated at 72 °C for 15 min for A tailing. The DNA was purified and cloned into the PMD19-T Simple vector (Takara, 6013). The cloned DNA was sequenced by the Sanger method. For each of SSAs, 10 clones were sequenced.

### 2.6. Preparation of NGS libraries with tagmented chromatins of various cells by using SALP

We performed SALP-seq with 100,000 GM12878, HeLa, HepG2, and 293T cells, and various numbers of HepG2 cells (100,000, 50,000, 10,000, 5,000, 2,500, and 500). Cells were collected by spinning at 500 g for 5 min at 4 °C and washed once with 50 μL of cold phosphate buffered solution (PBS). Cells were lysed by resuspended in cold lysis buffer (10 mM Tris-HCl, pH 7.4, 10 mM NaCl, 3 mM MgCl_2_ and 0.1% IGEPAL CA-630). Cells were then spun at 500g for 10 min at 4 °C to collect the nuclei precipitate. For tagmentation of 100,000 cells, nuclei were tagmented in a 30-μL reaction containing 20 μL of Tn5 transposome, 3 μL of DMF and 1× LM buffer. For tagmentation of 50,000 and 10,000 cells, nuclei were tagmented in a 30-μL reaction containing 4 μL of Tn5 transposome, 3 μL of DMF and 1× LM buffer. For tagmentation of 5,000, 2,500 and 500 cells, nuclei were tagmented in a 5-μL reaction containing 1 μL of Tn5 transposome, 0.5 μL DMF and 1× LM buffer. The tagmentation reactions were mixed gently and incubated at 37 °C for 30 min, during which the reactions were gently mixed per 10 min to improve the tagmentation efficiency. In tagmentation, different BTAs (Supplementary Table 1 and 2) were used to tagment the different cell samples. After tagmentation, the chromatins of 100,000 cells of 4 cell lines and 5 cell numbers of HepG2 cell were separately pooled together to generate two mixtures. To mixture, 1% SDS and 20 mg/mL Proteinase K (Sigma) were add to final concentrations of 0.1% and 400 μg/mL, respectively. The mixture was incubated at 65 °C for 1 hour and then added with 1× TE buffer to a final volume of 200 μL. The gDNA was then purified from the mixture by using standard phenol:chloroform extraction as described ^22^. For preparing NGS library, the purified gDNA was successively ligated with SSA, elongated with rTaq, and amplified with Illumina compatible index primers as described above. Finally, the DNA fragments of 150–1000 bp were recovered to include more nucleosome-free sequences. In library preparation, the SSA with 3N overhang was used according to SSA optimization.

### 2.7. Preparation of NGS libraries with Hind III-digested and sonicated DNA by using SALP

The Hind III-digested and sonicated gDNA was ligated with the SSA of 3N overhang and elongated with the same procedure as the tagmented DNA. Then the gDNA was ligated with a T adaptor in a 10-μL containing 1 μL of T adaptor (5 μM), 1× T4 DNA Ligase Reaction Buffer and 1 μL of T4 DNA ligase at 16 °C for 2 h. Finally, the gDNA was purified with 1.2× Ampure XP beads and amplified with different NEB index primers (Supplementary Table 1) in a 50-μL PCR reaction as described above. The PCR products were run with agarose gel and the gDNA fragments of 300–1000 bp were extracted with the QIAquick Gel Extraction Kit.

### 2.8. Next-generation sequencing

After amplified with Illumina-compatible primers (Supplementary Table 1), four NGS libraries, including NGS-L1 (tagmented chromatins of four cell lines), NGS-L2 (tagmented chromatins of five cell numbers), NGS-L3 (HindIII-digested gDNA), and NGS-L4 (sonicated gDNA), were constructed by using SALP. The libraries were detected and quantified with Agilent Bioanalyzer 2100. Four libraries were mixed together according to the DNA quality (ng) at the ratio of 4:1:1:1 (NGS-L1:NGS-L2:NGS-L3:NGS-L4) and sequenced by a lane of Illumina Hiseq X Ten platform (Nanjing Geneseeq).

### 2.9. Analysis of SALP-seq data

The raw reads data were separated according to the index and barcode by using a homemade Perl scripts. Then the ME (19 bp) and barcode (6 bp) sequences were removed from the 5′ end of the pair-end sequencing reads 2. All reads were cut to 30 bp and aligned to the human genome (hg19) by using Bowtie program (version 1.1.2),^23^, with the default settings, except that the parameter-X 2000 was used to ensure the long fragments could be aligned to the genome. Peak calling was performed with macs2,^24^, with the following parameters:-f BEDPE-keep-dup=2. Peak annotation was performed with Homer software.^25^ Gene ontology analysis was performed with PANTHER by uploading the genes to the website (http://pantherdb.org/).^26^ De novo motif analysis was performed with MEME tool and the comparison between enriched motifs and HOCOMOCO human v10 database was performed with Tomtom tool from MEME software suite.^27,28^ The BEDTools intersect was used to detect the overlapped peaks with –wa –u parameters.^29^ Microsatellites were predicted using MIcroSAtellite (MISA) script with the default parameters.^30^ All peak tracks were displayed with UCSC genome browser and statistical analysis were performed with R program and homemade Perl scripts. The ACAT-seq reads data of GM12878 was downloaded from the GEO database (accession number: GSE47753). The ACAT-seq reads data were analyzed as the SALP-seq reads data to compare.

### 2.10. Data availability

The raw reads data from SALP-seq are available at NCBI GEO with the accession number: GSE 104162.

## 3. Results

### 3.1. Development of tagmentation-based SALP method

The DNA fragments tagmented by Tn5 transposase have recently been used to facilely construct the NGS library. We firstly constructed the NGS library by using Tn5-tagmented DNA fragments with SALP method. For this end, a new kind of barcoded Tn5 adaptor (BTA) was designed (Figure 1A). The barcode can be used to differ the samples pooled together after the tagmentation. By using the complex of Tn5 transposase and BTA, the gDNA from HepG2 cells was efficiently fragmented (Supplementary Figure 1A), based on the “cut and paste” mechanism of Tn5.^31^ The tagmented fragments were then denatured and a single strand adaptor (SSA) were ligated. The SSA was a double-stranded oligonucleotide with a 3′ overhang of 1–4 random nucleotides. After ligation, the gDNA was elongated with Taq polymerase and amplified with PCR by using a pair of primers that can anneal with BTA and SSA. The PCR amplification reveals that the SSA with 3 base-overhang produced the highest ligation efficiency in all SSAs (Supplementary Figure 1B). To further confirm the constructed DNA library, the PCR products were cloned into T-vector and 40 clones identified by the colony PCR were sequenced by using Sanger method. The colony PCR detection indicated that the cloned DNA fragments had the size from 150 to 1000 bp (Supplementary Figure 1C). The sequencing revealed that the libraries were successfully constructed with 4 kinds of SSAs by using SALP method (Supplementary Figure 2), which were compatible with Illumina sequencing platform (Supplementary File 2). Based on these results, the SSA with 3-base overhang was subsequently employed to construct the following libraries.

**Figure 1.**
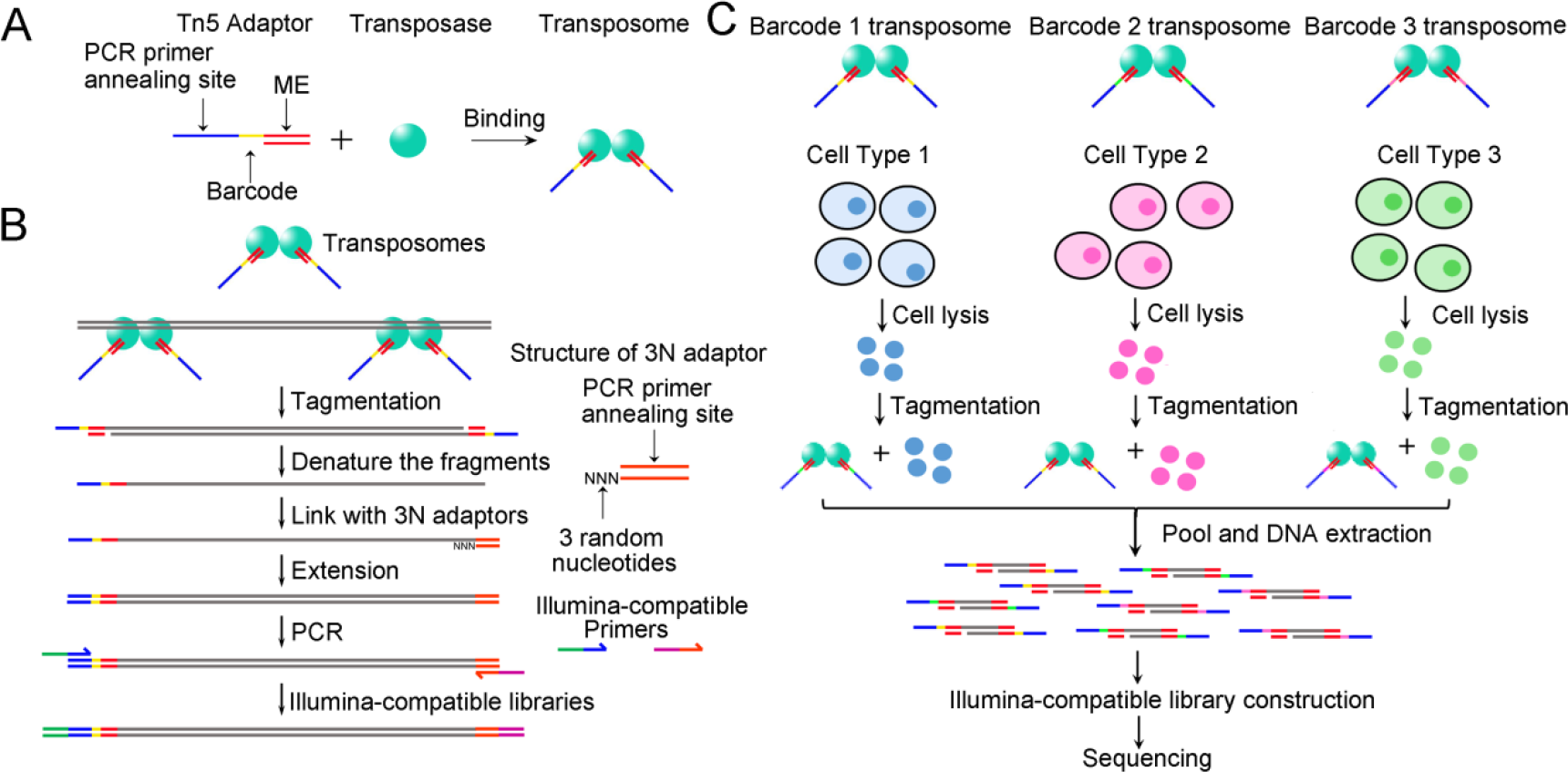
Schematic of tagmentation-based SALP-seq. (**A**) Barcoded Tn5 adaptor (BTA) used in SALP-seq. BTA consists of a 19-bp double-stranded transposase binding site (ME) and a single-stranded barcode and PCR primer annealing site. The Tn5·BTA complex (transposome) is used to tagment DNA or chromatin. (**B**) SALP-seq experimental procedures. Single strand adaptor (SSA) is a double-stranded oligonucleotide with a 3′ overhang, which is 3 random nucleotides (3N). (**C**) Tagmentation-based SALP-seq strategy for high-throughput characterization of chromatin openness states of multiple cells.

### 3.2. A high-throughput procedure for preparing tagmentation-based NGS library with SALP

Tn5 tagmentation has been recently used to profile the open chromatin regions in a technique named as ATAC-seq. Therefore, we then applied SALP in combination with NGS (SALP-seq) to the same field. Based on the BTA designed by this study, as many as 10 different BTAs were designed and used to prepare the NGS library for profiling the open chromatin regions in a facile high-throughput procedure (Figure 1B). Four different cell lines including GM12878, HepG2, HeLa and 293T were respectively tagmented with the complexes of Tn5 and BTAs with different barcodes. The tagmented chromatins of four cell lines were then pooled together. Similarly, five different numbers of HepG2 cells were also respectively tagmented with the complexes of Tn5 and BTAs with other different barcodes. The tagmented chromatins of five cell numbers were then pooled together. DNA was then extracted from the pooled chromatins and two NGS libraries were prepared with the SALP procedure verified above. The two NGS libraries, together with two NGS libraries prepared with the enzymatically and physically fragmented gDNAs by using SALP, were then mixed together and sequenced by a lane of Illumina Hiseq X Ten platform. As a result, as many as 95,202,715 mappable reads were obtained (Supplementary Table 3).

### 3.3. Characterization of chromatin openness state of lymphoblastoid with SALP-seq

The chromatin state of GM12878 cell had been widely explored by using various methods including DNase-seq, FAIRE-seq and ATAC-seq. We also explored the chromatin state of this cell line by using SALP-seq for comparing the SALP-seq results with those obtained by other methods. The genome-wide reads distribution of SALP-seq and ATAC-seq ^32^ was firstly compared, which revealed that the similar genome-wide reads distribution was obtained by two methods (Figure 2A). Some reads highly enriched regions were commonly identified by two methods. However, some reads highly enriched regions were only identified by SALP-seq or ATAC-seq. Comparison of peaks generated by two methods showed that SALP-seq found more peaks than ATAC-Seq according to the sequencing depth normalized peak numbers (Figure 2B). Calculation of reads density of co-located peaks revealed that the reads density of SALP-seq peaks were higher than that of ATAC-seq peaks (Figure 2C). Comparison of fold enrichment of peaks demonstrated that the peak with low fold enrichment could be more easily identified by SALP-seq (Figure 2D). These data indicate that SALP-seq could more sensitively find the open chromatin regions than ATAC-seq. The comparison of reads distribution demonstrated that the SALP-seq obtained the same reads density distribution as ATAC-seq (Figure 2E), indicating the reliability of SALP-seq. For further confirming the reliability of SALP-seq in characterizing the chromatin state, the peaks in a locus (Figure 2F) was compared to those previously highlighted by other methods,^33^ which revealed that the same chromatin openness state was obtained by SALP-seq, ATAC-seq, FAIRE and DNase-seq. In addition, the SALP-seq peaks are well matched with the H3K27Ac track peaks and DNase clusters, further indicating the reliability of SALP-seq in characterizing the chromatin state.

**Figure 2.**
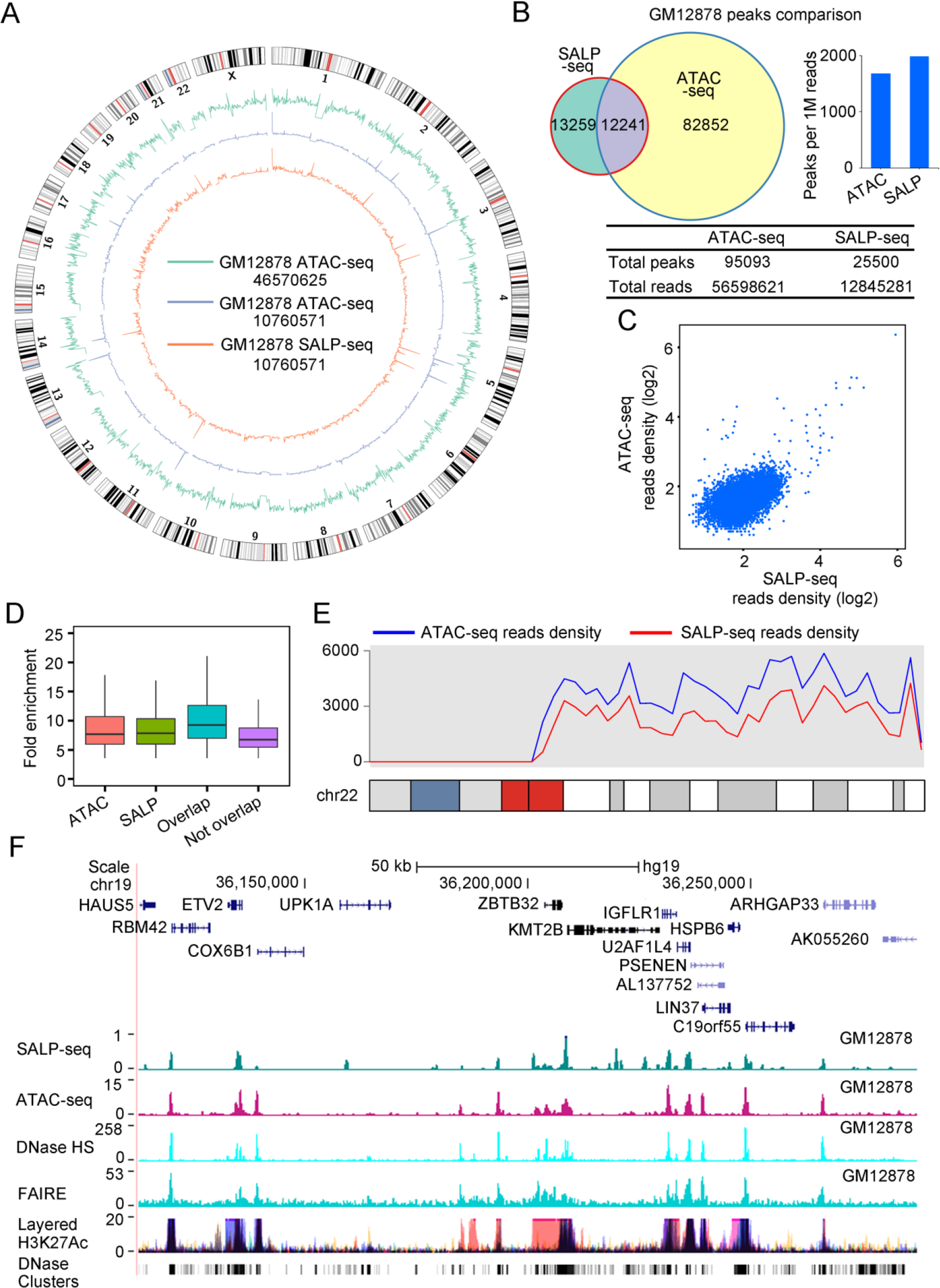
Comparative characterization of chromatin openness states of GM12878 cell line by SALP-seq and ATAC-seq. (**A**) Distribution of reads density. The reads density refers to the reads number in each 1-Mb window. (**B**) Comparison of peaks identified by two methods. (**C**) Comparison of reads density of common peaks identified by two methods. (**D**) Fold enrichment of various peaks. ATAC, ATAC-seq peaks. SALP, SALP-seq peaks; Overlap, Overlapped ATAC-seq and SALP-seq peaks; Not Overlap, SALP-seq peaks not overlapped with SALP-seq peaks. (**E**) Distribution of reads density on chromosome 22. (**F**) UCSC browser track showing chromatin openness state of a selected region characterized by SALP-seq and other methods.

### 3.4. Characterization of chromatin openness states of different cells with SALP-seq

Based on BTAs designed by this study, a NGS library was prepared with four cell lines including GM12878, HepG2, HeLa and 293T in a high-throughput approach by using SALP (Figure 1B). The calculation of reads density around of transcription start sites (TSSs) revealed that the TSS regions showed the high reads density (Figure 3A), indicating the high-level chromatin openness in these regions as previously reported ^34^. For comparing the chromatin openness states among different cell lines, reads density were calculated in the whole genome scale (Figure 3B). It is clear that some regions showed as the common open chromatin regions in all cell lines, such as regions located in chromosome 5. As an example, a chromatin locus in chromosome 19 were shown, which revealed that the highly enriched SALP-seq peaks existed in all cell lines and agreed with the H3K27Ac sites and DNase clusters generated by ENCODE Consortium (Figure 3C). At the genome-wide scale, many co-located peaks were found in different cell lines, indicating that there are many common chromatin open regions among different cells. However, some common chromatin open regions showed the different open level. Additionally, each cell line contains its specific peaks, indicating the cell-type-specific chromatin openness states among different cells. These data reveal that SALP-seq provides an easy method to perform the comparative characterization of chromatin states of different cells.

**Figure 3.**
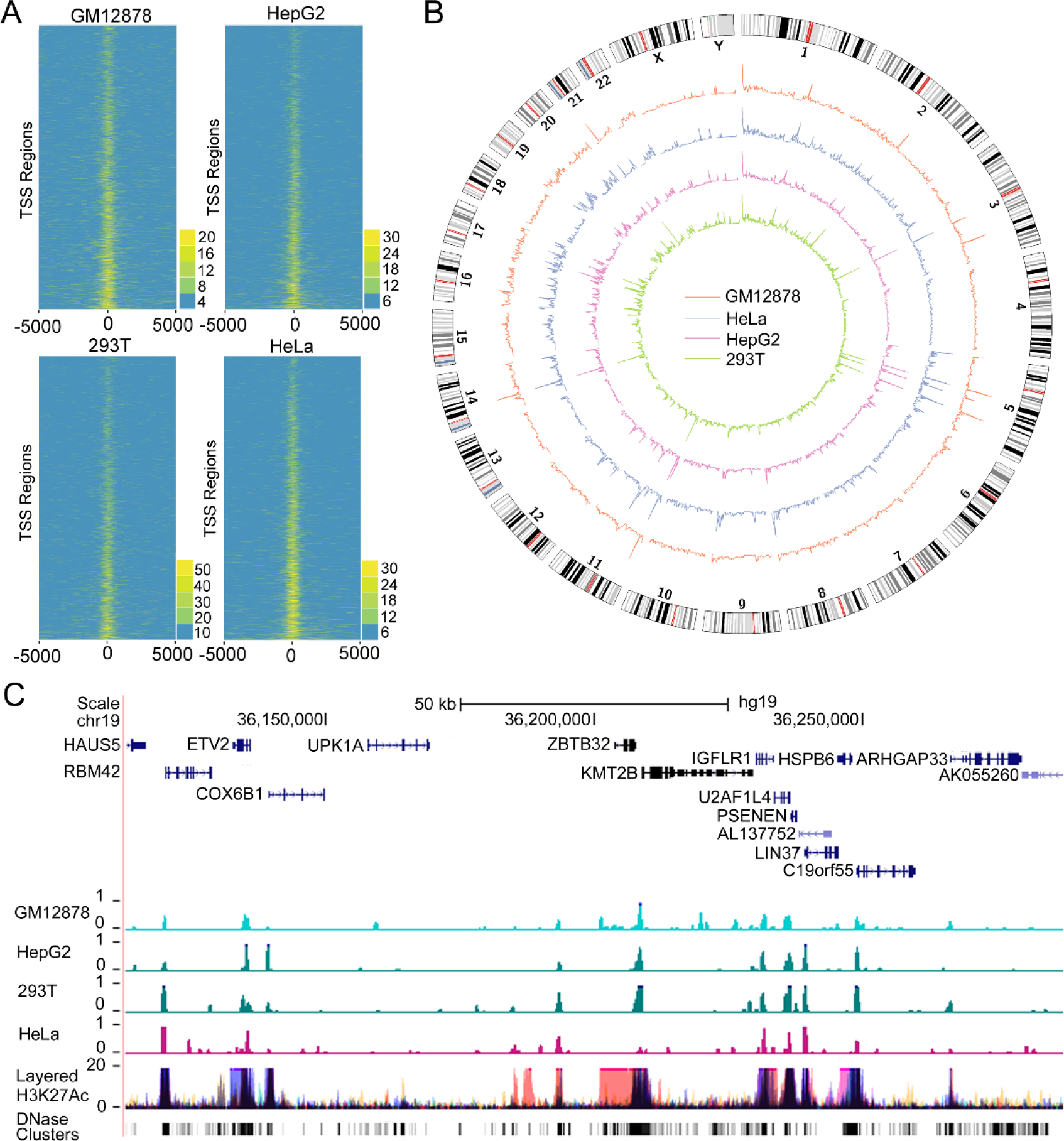
Comparative characterization of chromatin openness state of four cell lines by SALP-seq. (**A**) Reads distribution around TSS. (**B**) Distribution of reads density. Reads density refers to the reads number in each 1-Mb window. (**C**) UCSC browser track showing the SALP-seq peaks in a selected genome locus. The chromatin openness indicators, layered H2K27Ac and DNase Clusters, of ENCODE were also shown.

### 3.5. Characterization of open chromatin regions with different number of cells by SALP-seq

As the methods to detect the chromatin openness, FAIRE-seq,^35^ and DNase-seq,^36^ usually requires 1–50 million cells, but ATAC-seq requires 50000 to 500 cells.^32^ To validate that the chromatin accessible regions could be identified with different numbers of cells by using SALP-seq, we performed SALP-seq with different numbers of HepG2 cells, including 100,000, 50,000, 10,000, 5,000, 2,500, and 500 cells. The reads density of each number of cells was calculated on genome scale, which revealed that the chromatin regions with high accessibility were commonly identified by 5 different numbers of cells (Figure 4A and 4B). The open chromatin regions identified by SALP-seq are matched with the H3K27ac track and DNase Clusters of ENCODE (Figure 4B). Nevertheless, the sensitivity was diminished when low number of cells were used as starting material (Figure 4C). With low numbers of cells, only most of open chromatin regions with high FE were identified ((Figure 4D). It means that SALP-seq can identify those highly open chromatin regions even with as few as 500 cells.

**Figure 4.**
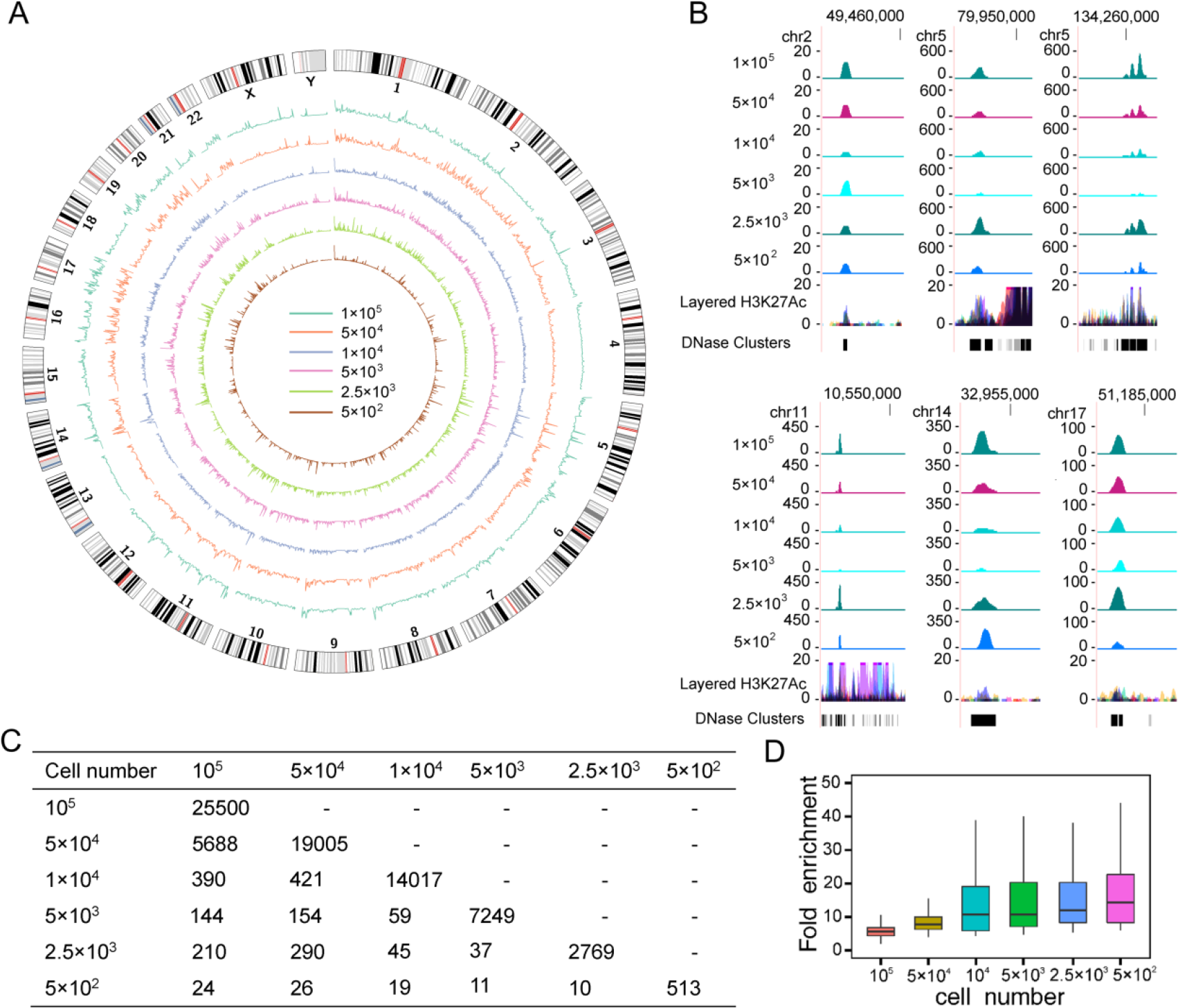
Chromatin openness state captured by SALP-seq with different numbers of HepG2 cells. (**A**) Reads density of different numbers of cells in whole genome scale. (**B**) UCSC browser track showing some typical open chromatin regions identified with different numbers of cells by SALP-seq. The chromatin openness state identified by ENCODE was also shown to compare. **(C)** Peaks obtained with different numbers of cells. The overlapped peaks were shown. **(D)** Fold enrichment of peak obtained with different numbers of cells.

### 3.6. Construction of NGS library with enzymatically digested or sonicated gDNA by SALP-seq

To show that SLAP could be used to construct NGS library with any other DNA fragments, we finally constructed NGS libraries with Hind III-digested and sonicated HepG2 gDNA by using an adapted SALP procedure (Figure 5A). In this procedure, after the SSA was ligated and elongated, a T adaptor with one T base overhang was ligated to the other end of DNA fragments. Then DNA was amplified with two primers annealing to SSA and T adaptor for generating the Illumina compatible sequencing libraries. The results revealed that the NGS libraries could be successfully constructed with the enzymatically and physically sheared gDNAs (Supplementary Figure 3), which were also sequenced by using Illumina Hiseq platform together with tagmented DNA. For checking the coverage of the constructed libraries, the genome-wide reads density of two NGS libraries were calculated. The results indicated that the reads were uniformly distributed on chromosomes (Figure 5B and 5C; Supplementary Figure 4 and 5), indicating that the NGS library could be successfully constructed with the enzymatically and physically sheared gDNAs by using SLAP. Additionally, the genome-wide distributions of the predicted Hind III digestion sites basically matched with the reads density of the NGS library constructed with Hind III-digested gDNA (Figure 5B and Supplementary Figure 4). However, some significant difference were also found in several chromosomes such as Chromosome 5, 9 and 11 (Supplementary Figure 4), which may be caused by the difference in sequences between the reference genome (hg19) and the HepG2 cell genome.

**Figure 5.**
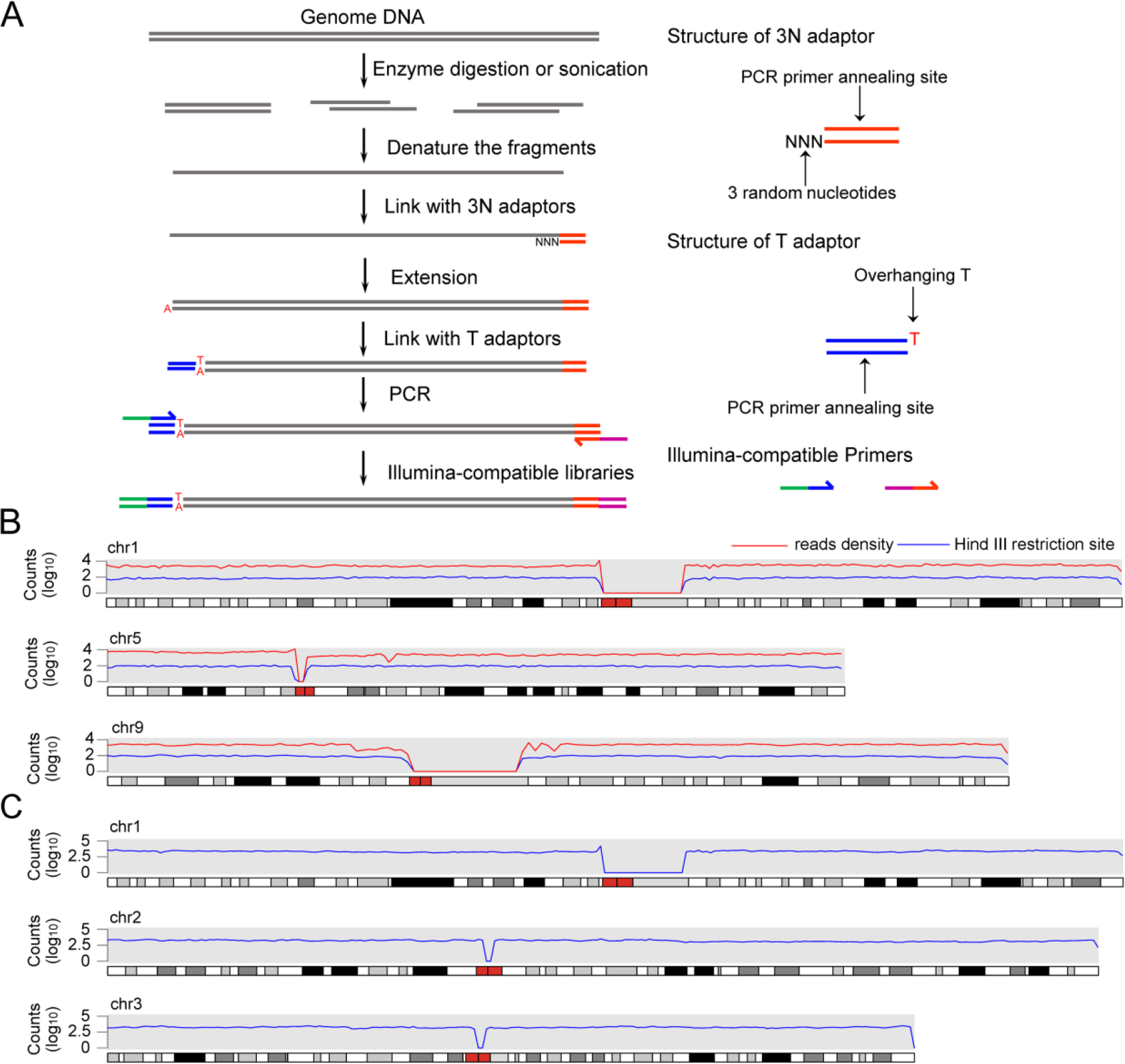
NGS library construction with alternatively sheared gDNA by using SALP method. (**A**) Schematic of library construction with the enzymatically digested and sonicated DNA by using SALP method. (**B**) Distribution of reads density of NGS library constructed with the Hind III-digested HepG2 gDNA and the density of predicted Hind III restriction sites in whole genome. (**C**) Distribution of reads density of NGS library constructed with the sonicated HepG2 gDNA. The information of other chromosomes was shown as supplementary Figure 5 and 6.

## 4. Discussion

In the current NGS era, as development of the next-generation sequencing, inexpensive production of large volumes of sequence data has great advantage over conventional methods ^2^. With this advantage, NGS has been used in virtually all branches of biological and medical research. As the fundamental step of NGS, the library construction methodology has been attracted increasing attention. Here, we report the development of a novel method, named as SALP, for constructing NGS library. SALP can be used to construct high-quality NGS libraries with enzymatically or physically sheared gDNA. Importantly, SALP can be used to construct high-quality NGS libraries based on Tn5 tagmentation in an unbiased high-throughput format, which can be widely used in the current increasing chromatin state research.

With the feature of “cut and paste” of the transposase, a Tn5-based library construction method, named ATAC-seq, has significant advantages over conventional library construction methods, such as fewer steps and lower input requirements. However, ATAC-seq is limited by an inherent drawback that as many as 50 percent of tagmented fragments could not be sequenced due to using two different Tn5 adaptors. The Tn5-based SALP overcomes this limitation. In the Tn5-based SALP procedure, only one kind of Tn5 transposome adaptor was used to assemble with Tn5. By using SSA, all fragments generated by Tn5 tagmentation can be sequenced. We also optimized the other key adaptor, SSA, for SALP. The SSAs with 3′ end overhang ranging from 1 to 4 random base were tested (Supplementary Figure 1), which demonstrated that the SSA with overhang of 3 random bases showed the highest ligation efficiency in the key step of SALP. Nevertheless, we found that the SSAs with overhang of 1–4 random bases all successfully produced library. The SSAs with overhang of 1–2 random bases showed limited ligation efficiency, which may be related to the reason that the overhangs of these SSAs might be too short to form the stable structure under the ligase reaction condition and few SSAs were linked to DNA fragments. The SSA with overhang of 3 random bases could anneal with the end of the single-stranded fragments efficiently and tightly, which allows ligase to ligate SSA efficiently. When the SSA with overhang of 4 random bases was used, stable hybrids could be formed by 3 of 4 random bases; however, a one-base gap or mismatch can be easily generated in the annealed hybrids, which can reduce the efficiency of ligation and elongation.

Chromatin organization with cell type-specificity can affect accessibility and activity of regulatory elements and the manifestation of unique cellular phenotypes.^37^ Chromatin openness state as an important feature of chromatin structure exerts significant influence on gene transcription. SALP-seq can be used to identify the open chromatin region easily. Comparing the open chromatin regions identified by SALP-seq and ATAC-seq of GM12878 cell line revealed that the reads density of SALP-seq were higher than that of ATAC-seq in co-located peaks (Figure 2C). This phenomenon is caused by the principle of SALP method. In SALP, a double-strand DNA (dsDNA) fragment was firstly denatured into two single-strand DNAs (ssDNAs), which were then linked with SSA and elongated into two dsDNAs like DNA replication. This process allows one input dsDNA fragment to be sequenced twice in SALP-seq, which can reduce the sequencing error and mutation robustly. This process allows SALP-seq to detect the low- or ultra-low-frequency mutations with strand specificity, which are often related to somatic mutations.^38^ Compared with ATAC-seq, SALP-seq can identify more peaks due to its capability to find peaks with low fold enrichment (Figure 2D). These results are caused by the fact that all Tn5 tagmented fragments could be sequenced in SALP-seq and many regions with lower openness state can thus be captured by SALP-seq but may be missed by ATAC-seq (Figure 2D).

The Tn5 adaptor used in SALP-seq are specially designed, which has a unique barcode sequence between the ME and PCR primer annealing sites (Figure 1). With this design, chromatins of different cell samples tagmented by different BTAs can be pooled together. The chromatin cocktail can then be treated as one DNA sample, which greatly simplifies the library construction of multiple cell or DNA samples. The chromatin openness state of various cells can be easily performed with minimal bias caused by library construction process, which is beneficial to the comparative characterization of chromatin state. This study compared the chromatin openness state of four difference cell lines including GM12878, HepG2, HeLa, and 293T with SALP-seq in this way, which clearly revealed the common and cell-type-specific open chromatin regions among 4 cell lines. In our knowledge, this is the first time to explore the chromatin state of different cells like this, which is currently realized by using multiple PCR amplifications of different cell samples with different Illumina index primers in ATAC-seq. The tagmentation-based SALP-seq should be useful for comparative characterization of chromatin state of clinical tissues.^21^

It should be pointed out that the NGS library construction strategy described in Figure 5A can also be used to construct DNA library in high throughput by using the barcoded SSA. In this case, a double-stranded barcode sequence can be placed between the 3′ overhang and primer-annealing site. By using such barcoded SSAs, the SSA-ligated ssDNAs from different DNA samples can be pooled together. The pooled DNAs can be treated as one DNA sample to undergo the followed library construction steps, including elongation, T adaptor ligation and PCR amplification. As using the barcoded Tn5 adaptors, this can simplify the experimental operation, reducing artificial bias, and saving the labor, time and cost.

Finally, SALP should be useful for the NGS library construction of highly degraded DNA and free blood DNA, including circulating free DNA (cfDNA), circulating tumor DNA (ctDNA), and cell-free fetal DNA (cffDNA). Due to the high SSA ligation efficiency, SALP should be useful for the amplification, detection, and NGS library construction of free blood DNA and highly degraded DNA. SALP should find its wide applications in liquid biopsy, in vitro diagnostic products (IVD), non-invasive prenatal testing (NIPT), forensic medicine, and even paleontology and archaeology (treating ancient DNA) in the future. Several single-stranded DNA library preparation methods have been already used in these fields.^5–8^ However, the wide applications of these methods are still limited by their intrinsic drawbacks. In our opinion, SALP overcomes these shortcomings due to its simplicity and high efficiency.

In conclusion, we have developed a new method, named SALP, for constructing NGS library, which is based a novel single strand adaptor linking technique. This method can be used to construct NGS library with the DNA fragments sheared by various methods including tagmentation, sonication and enzymatic digestion. This method constructs NGS library in several simple steps just dependent on two cheap routine enzymes, T4 DNA ligase and Taq polymerase. Combining with Tn5 tagmentation technique, this method can simply construct NGS library in five steps, including tagmentation, SSA ligation, elongation, T adaptor ligation, and index PCR amplification. The Tn5-based SALP-seq overcomes the key shortcoming of the current ATAC-seq technique and develops a high-throughput procedure for constructing NGS libraries with multiple cell samples. The Tn5-based SALP-seq can thus be widely used to characterize the comparative chromatin openness states or genome sequences of various tissues or cells, such as cancerous tissues from various patients, which may be contribute to the current increasing personalized and precision medicine. SALP can construct NGS library with low number of cells in a high-throughput format, which is also useful for analyzing clinical tissue samples.

## Acknowledgements

This work was supported by the National Natural Science Foundation of China (61571119).

## Accession numbers

GSE 104162

## Data availability

The raw reads data from SALP-seq are available at NCBI GEO with the accession number: GSE 104162.

## Author contributions

J.K.W. conceived the idea and designed and instructed the study. J.W. performed main laboratory experiments and analyzed the data. W.D. prepared cells. L.W. prepared some library. J.W. and J.K.W. and wrote the paper.

## Supplementary data

Supplementary Data are available at Online.

## Supplementary information

**Supplementary Table 1.**
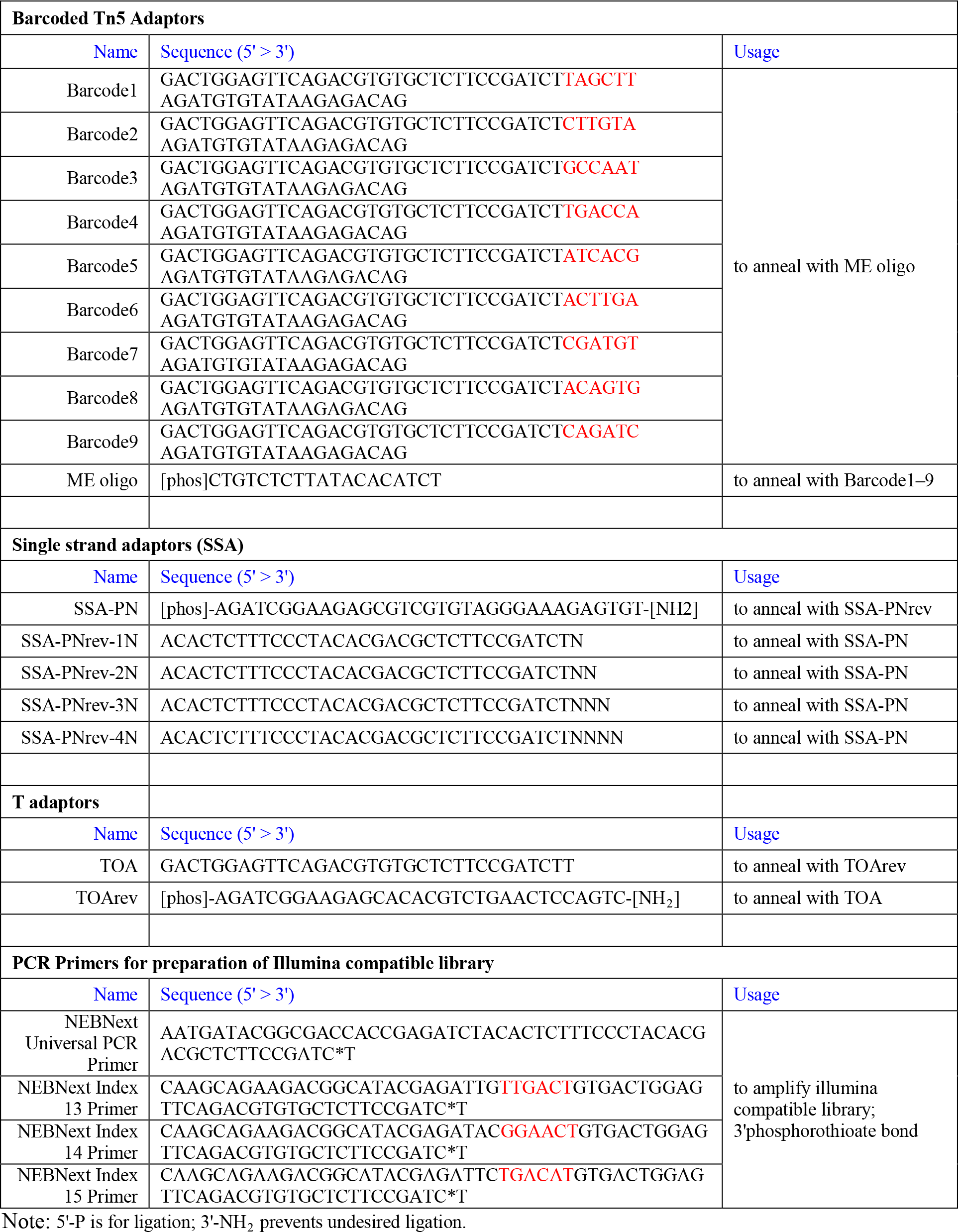
Oligonucleotides used as adaptors and PCR primers

**Supplementary Table 2.**
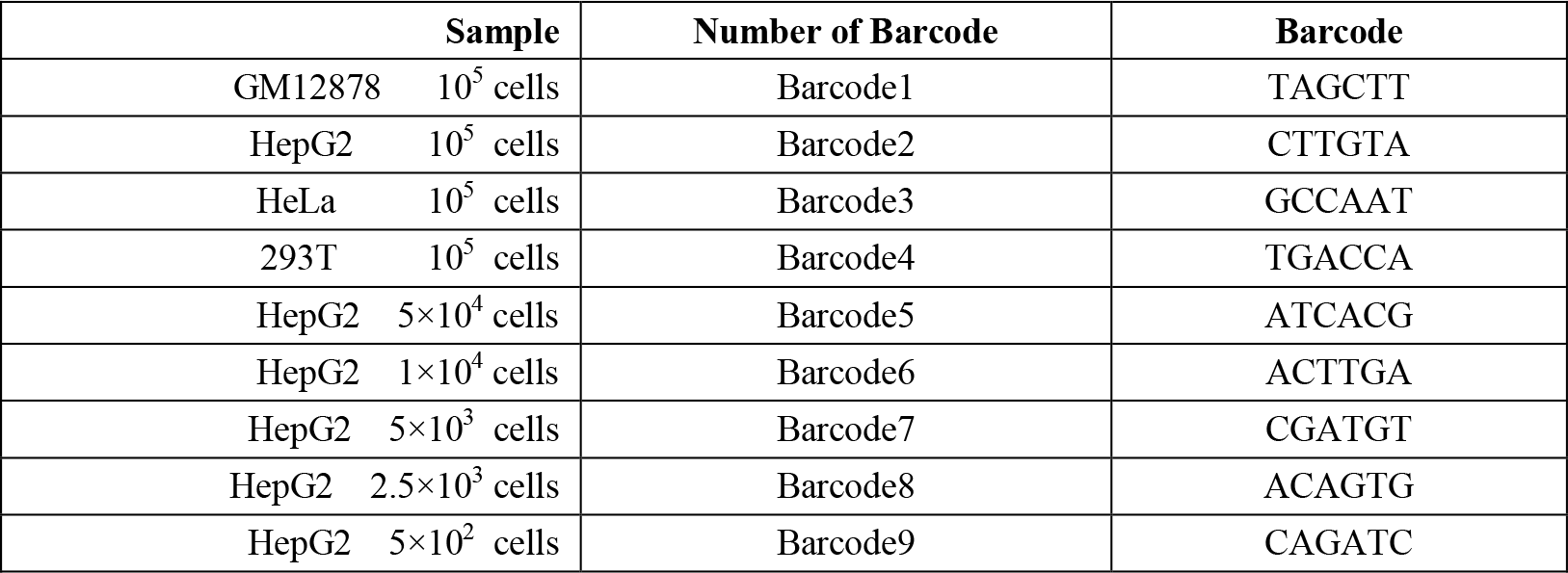
Barcodes on Barcoded Tn5 adaptors for labeling different cell samples.

**Supplementary Table 3.** Reads from a lane of Illumina Hiseq X Ten sequencing.

| Pooled DNA samples | Barcode | Index | NGS library | Reads number | Mappable reads number |
| --- | --- | --- | --- | --- | --- |
| GM12878 10<sup>5</sup> cells | TAGCTT | AGTCAA | NGS-L1 | 12845281 | 10760571 |
| HepG2 10<sup>5</sup> cells | CTTGTA |  |  | 17474263 | 12329768 |
| HeLa 10<sup>5</sup> cells | GCCAAT |  |  | 19120478 | 16439189 |
| 293T 10<sup>5</sup> cells | TGACCA |  |  | 10647259 | 8828634 |
| HepG2 5×10<sup>4</sup> cells | ATCACG | AGTCAA | NGS-L2 | 27245787 | 20946874 |
| HepG2 1×10<sup>4</sup> cells | ACTTGA |  |  | 399898 | 212125 |
| HepG2 5×10<sup>3</sup> cells | CGATGT |  |  | 134725 | 95232 |
| HepG2 2.5×10<sup>3</sup> cells | ACAGTG |  |  | 116644 | 86011 |
| HepG2 5×10<sup>2</sup> cells | CAGATC |  |  | 26425 | 7356 |
| HindIII-digested HepG2 gDNA | - | ATGTCA | NGS-L3 | 32944329 | 14140187 |
| Sonicated HepG2 gDNA | - | AGTTCC | NGS-L4 | 51687071 | 11356768 |

## Supplementary figures

**Supplementary Figure 1.**
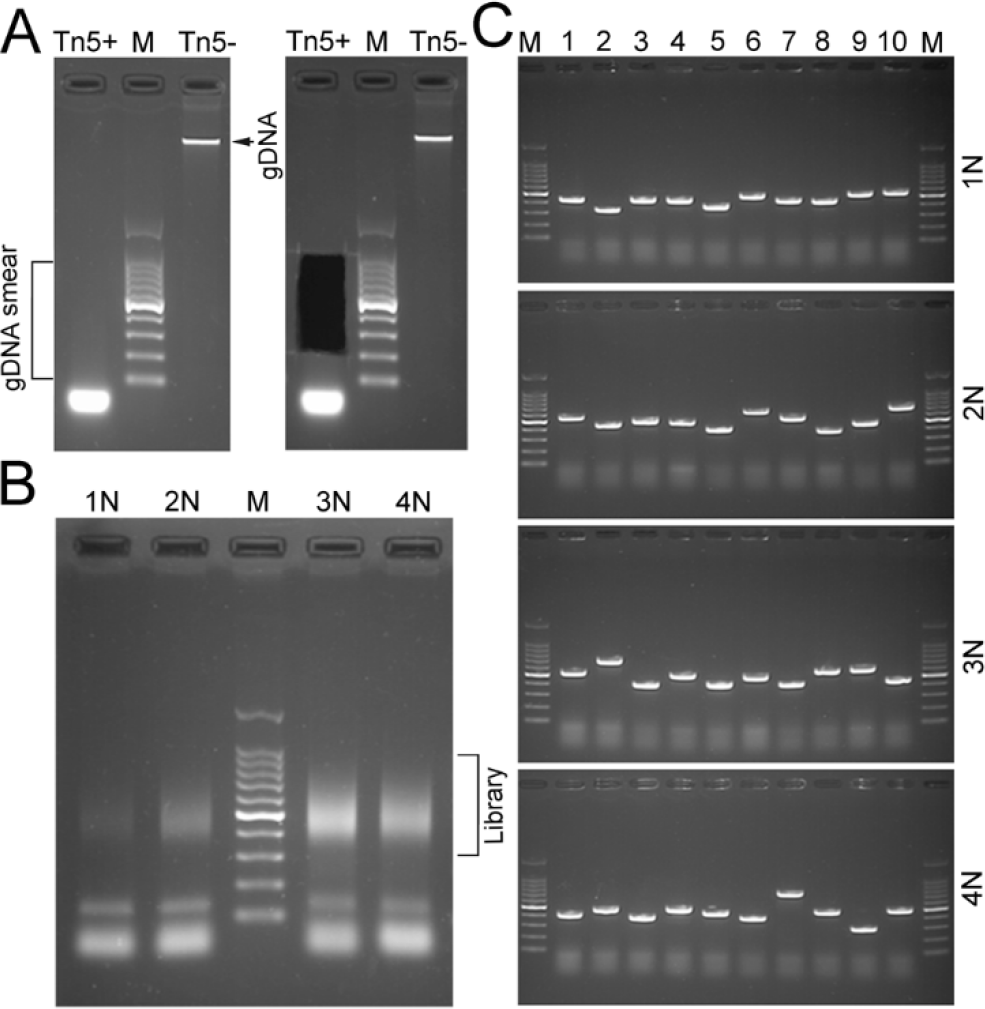
Validation of SALP method. (**A**) The HepG2 genomic DNA (gDNA) tagmented by Tn5 transposome. The tagmented gDNA showed as smear in comparison with the input DNA. DNA smear was cut and purified. (**B**) Illumina compatible libraries were constructed using different SSAs with different numbers of overhanging bases at 3′ end. The SSA with 3N overhang was adopted for its high-ligation efficiency. 1N–4N: SSAs with different numbers of random bases. (**C**) Clone sequencing was performed with the libraries constructed with 4 different SSAs to check the libraries structure. From top to bottom: colons from 1N to 4N libraries.

**Supplementary Figure 2.**
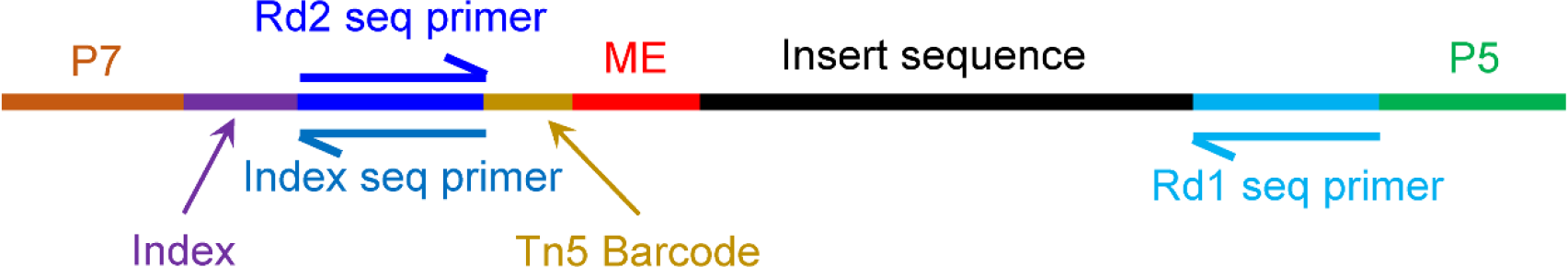
The structure of SALP library. The SALP sequencing library structure compatible with Illumina sequencing platform was illustrated. And the name of the each element were shown in the figure.

**Supplementary Figure 3.**
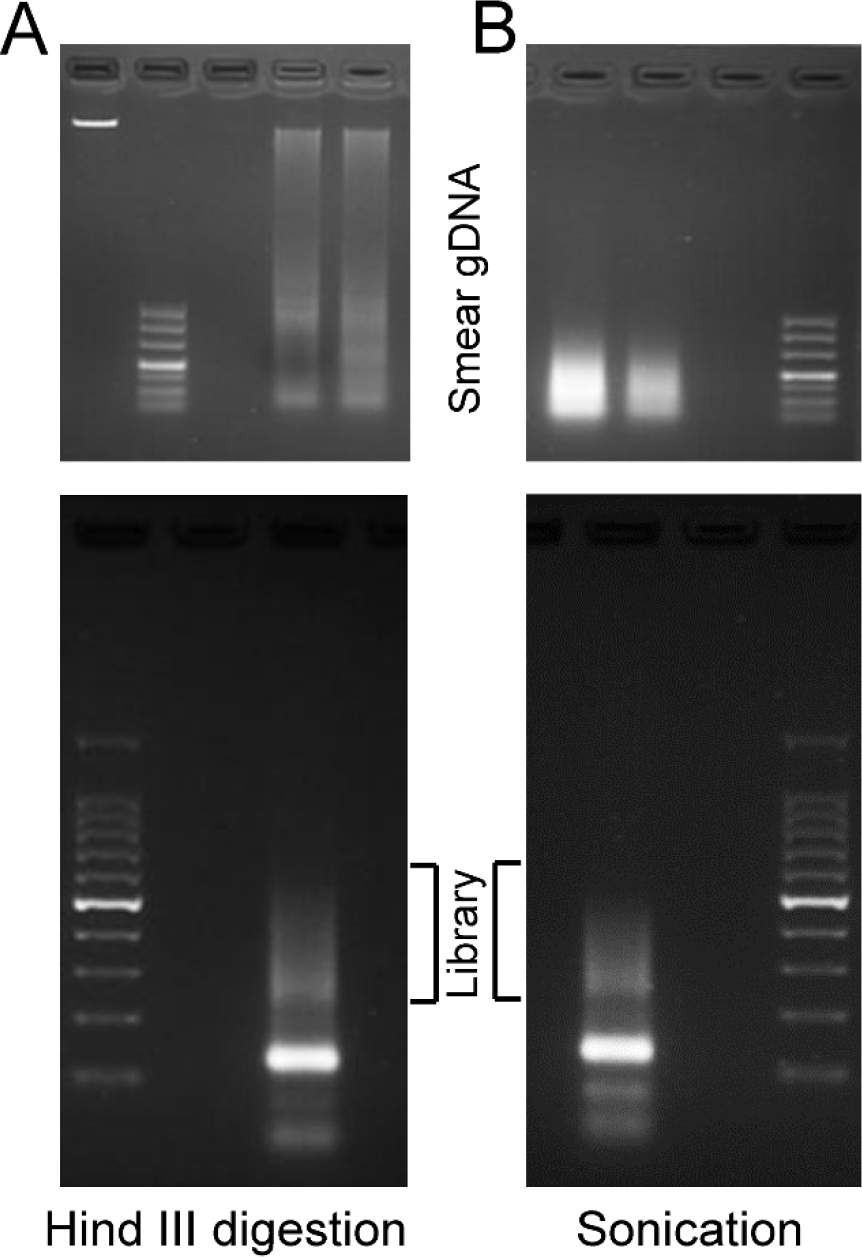
The sequencing libraries constructed by SALP started with different fragmentation methods. (**A**) The library constructed with the Hind III-digested HepG2 genomic DNA. The HepG2 genomic DNA digested by Hind III were shown on the top, and constructed library were shown on the bottom. (**B**) The library constructed with the sonicated HepG2 genomic DNA. The HepG2 genomic DNA sheared with sonication was shown on the top, and the library constructed with sonicated DNA was shown on the bottom.

**Supplementary Figure 4.**
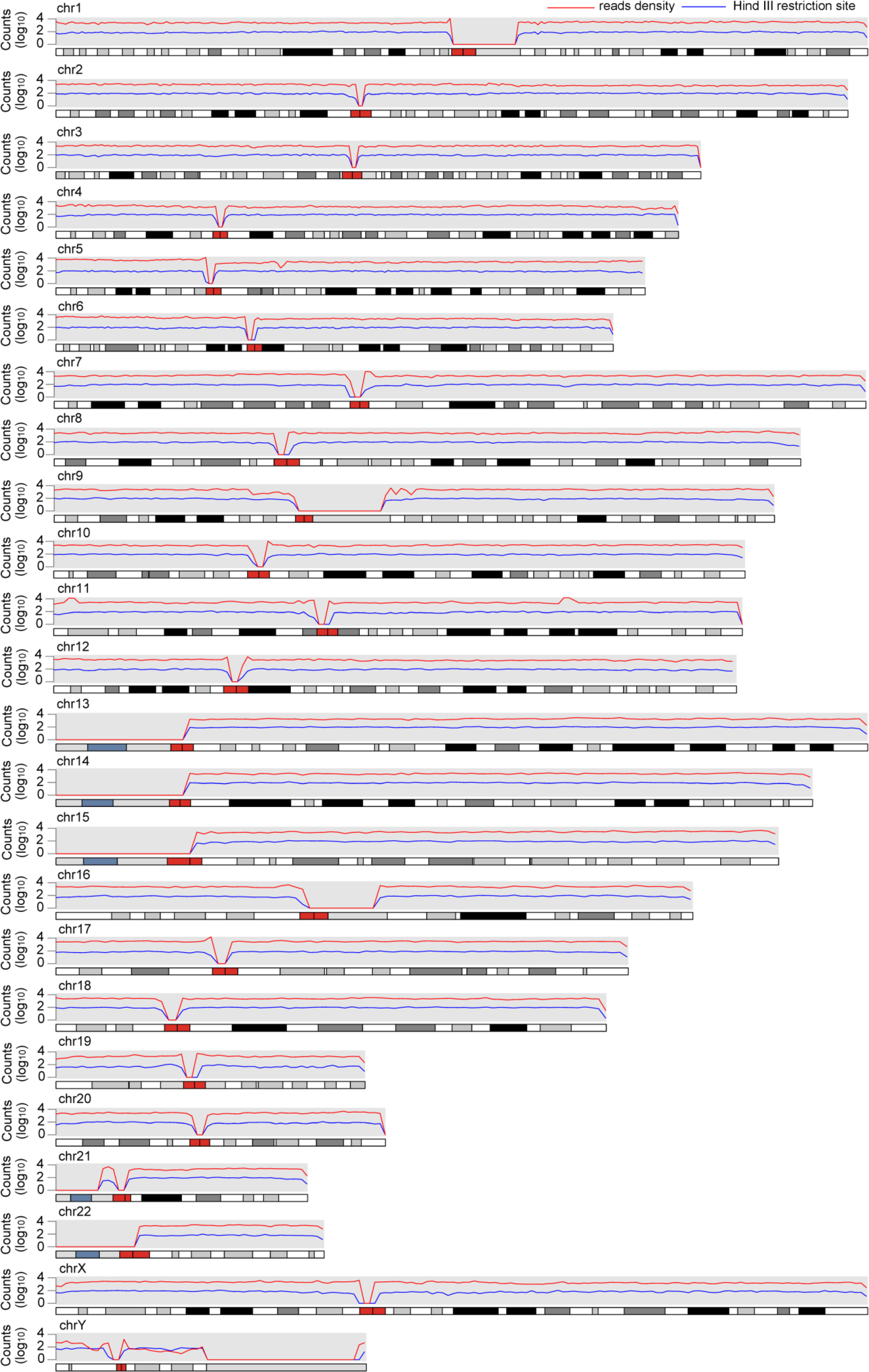
Comparison of the distribution of Hind III digestion library reads density and Hind III restriction sites through the whole genome. The Hind III digestion library reads density and Hind III restriction sites density were calculated through the whole genome scale with 1M window.

**Supplementary Figure 5.**
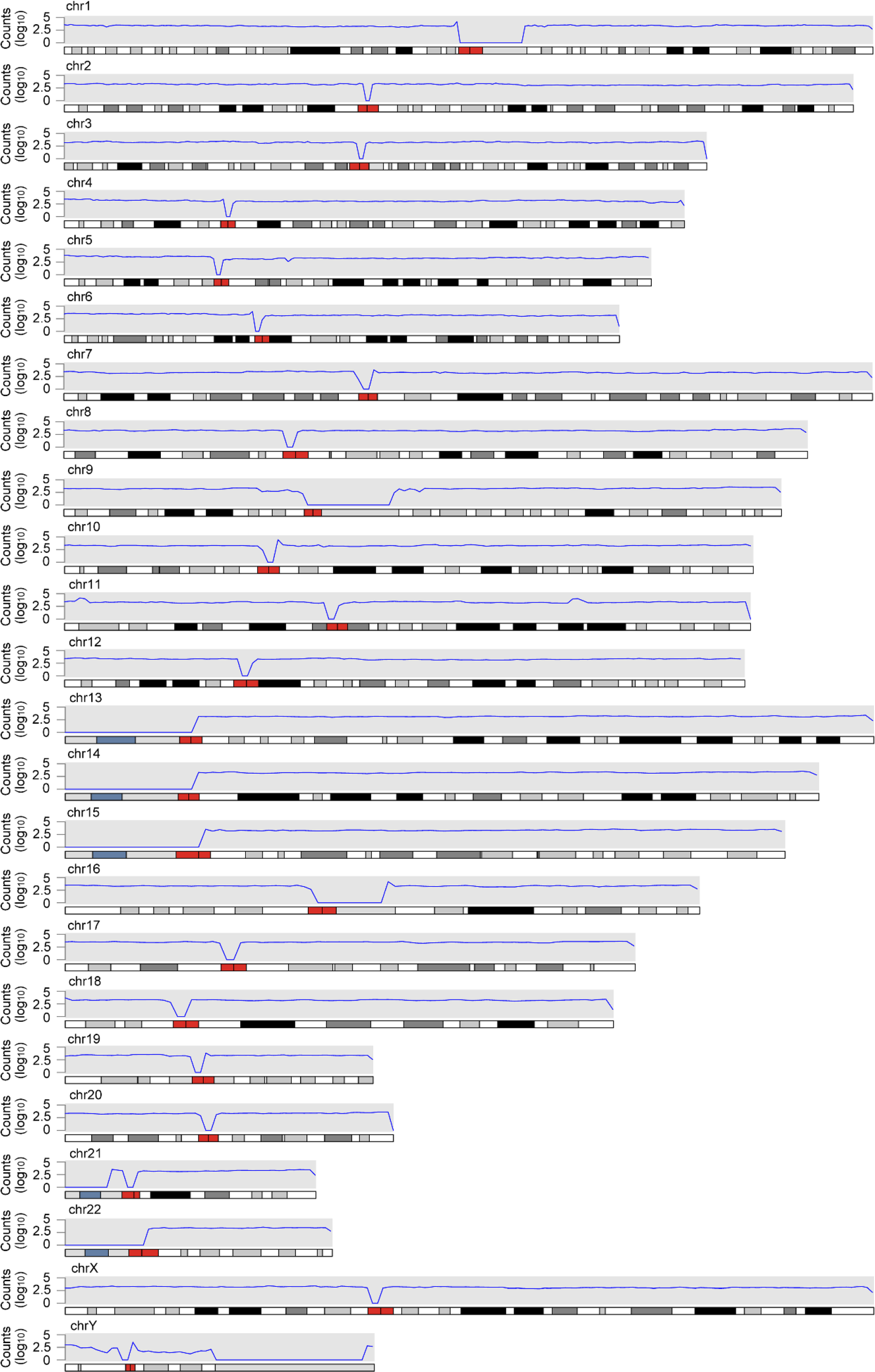
The reads distribution of sonication library. The reads dense of sonication library in whole genome scale were calculated with 1 M window.

## Supplementary File 2

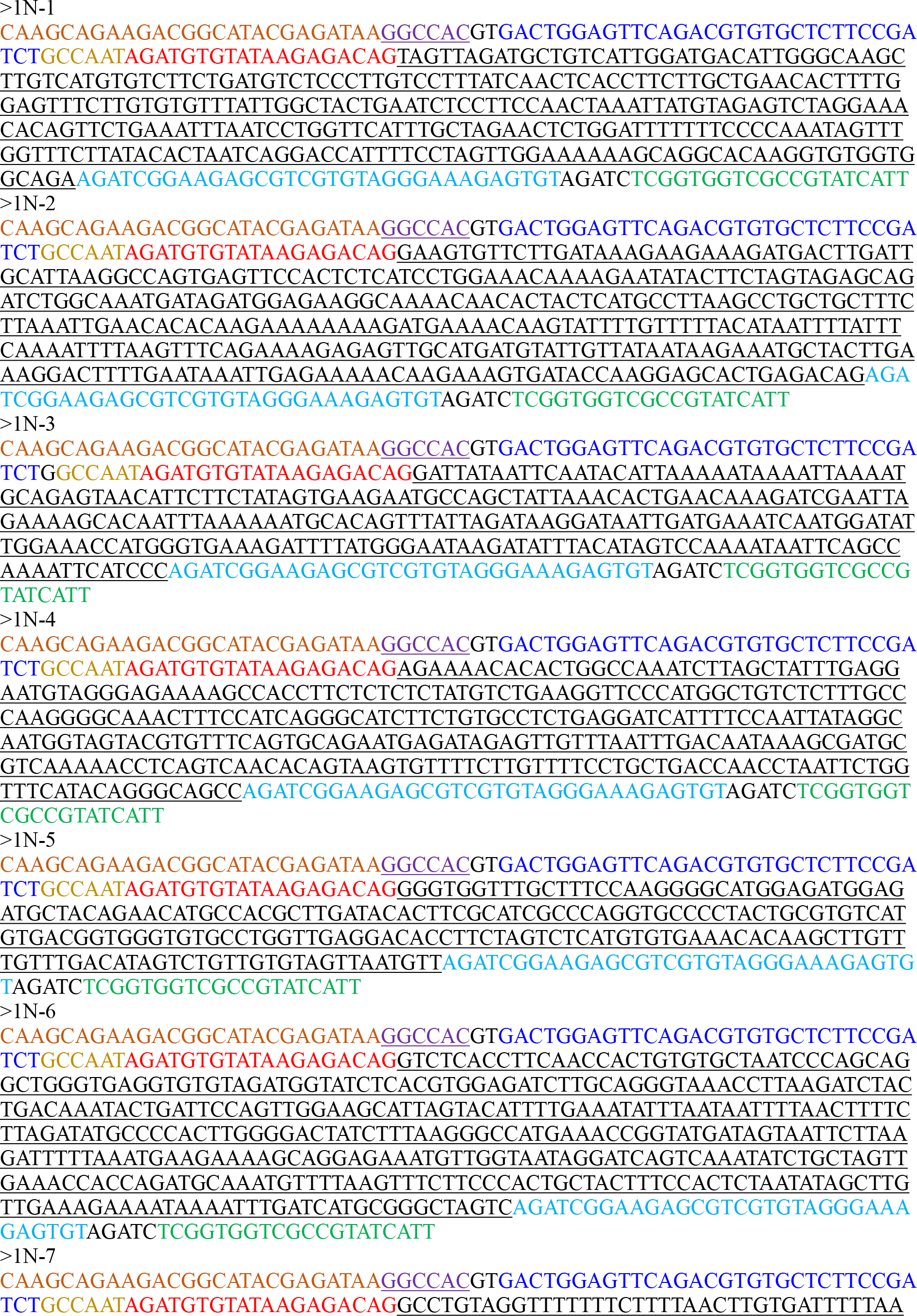

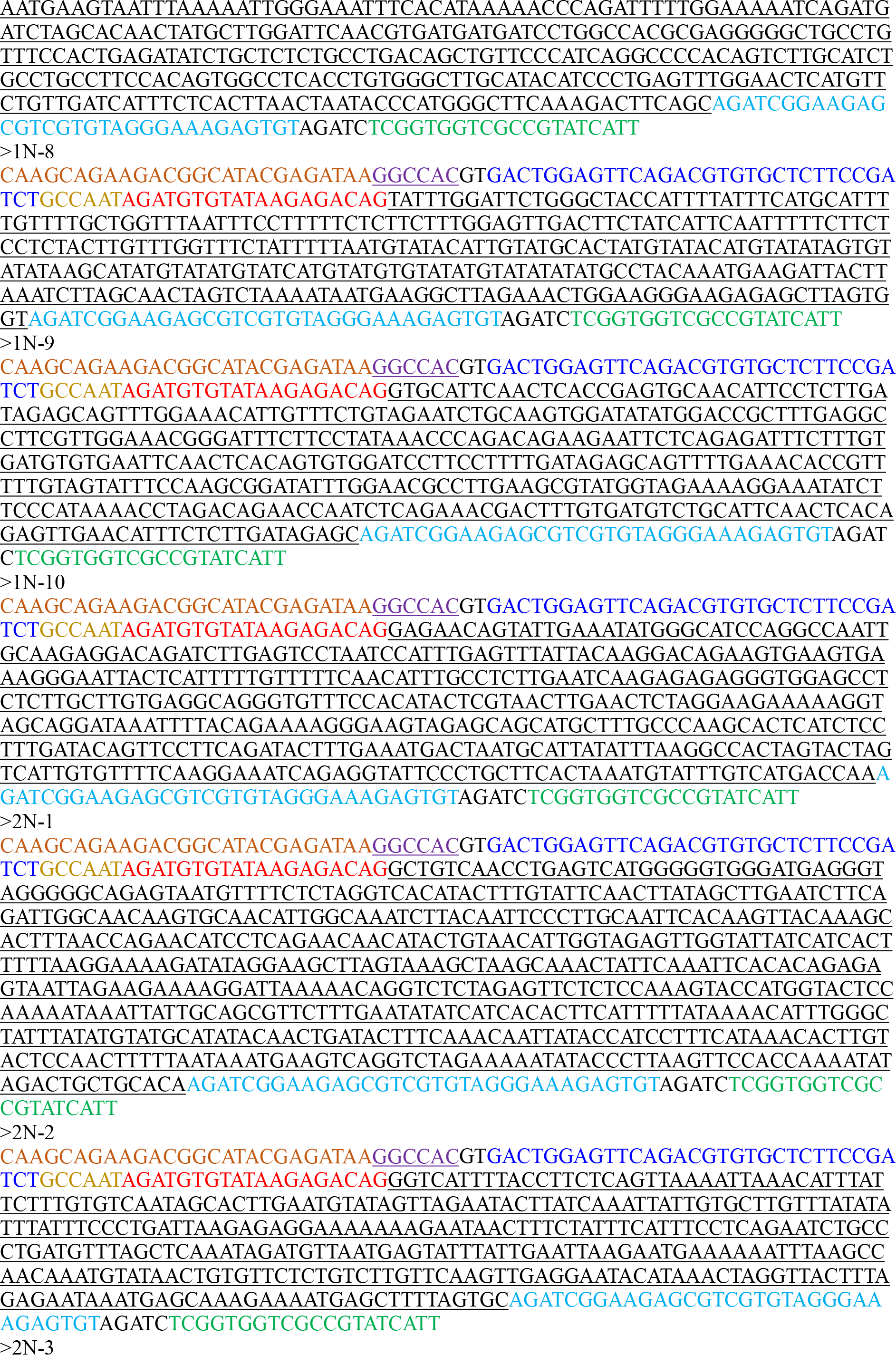

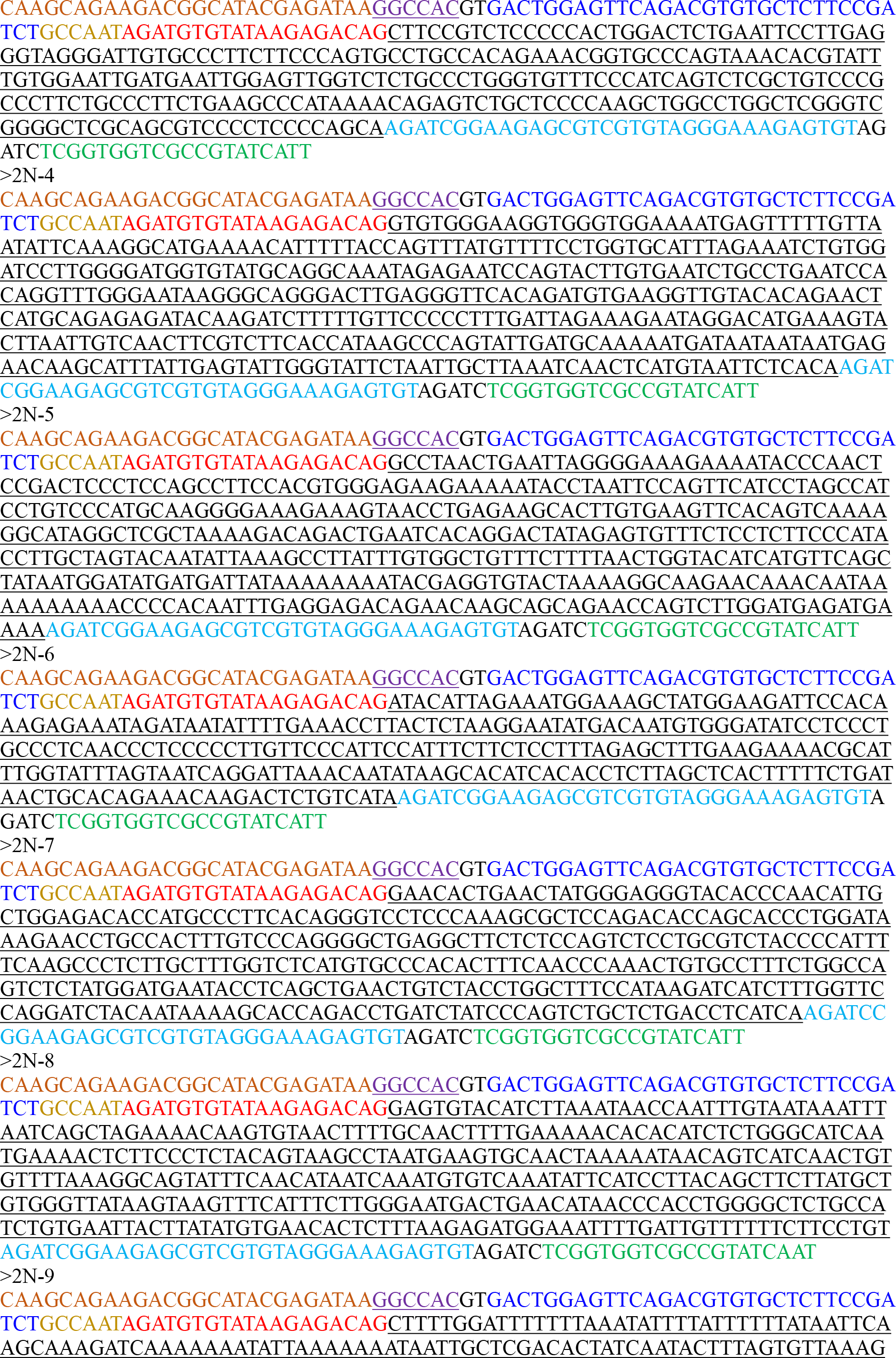

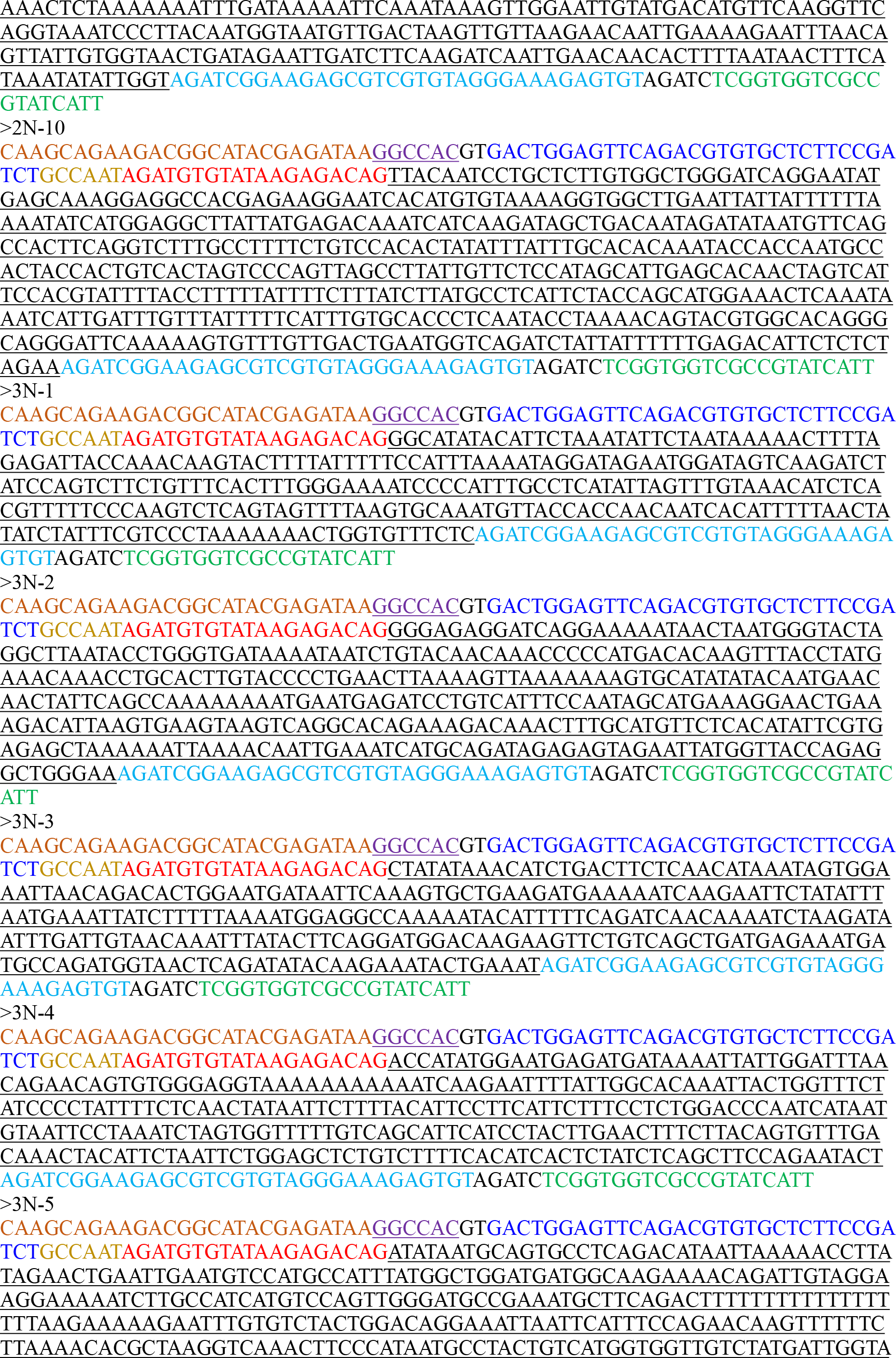

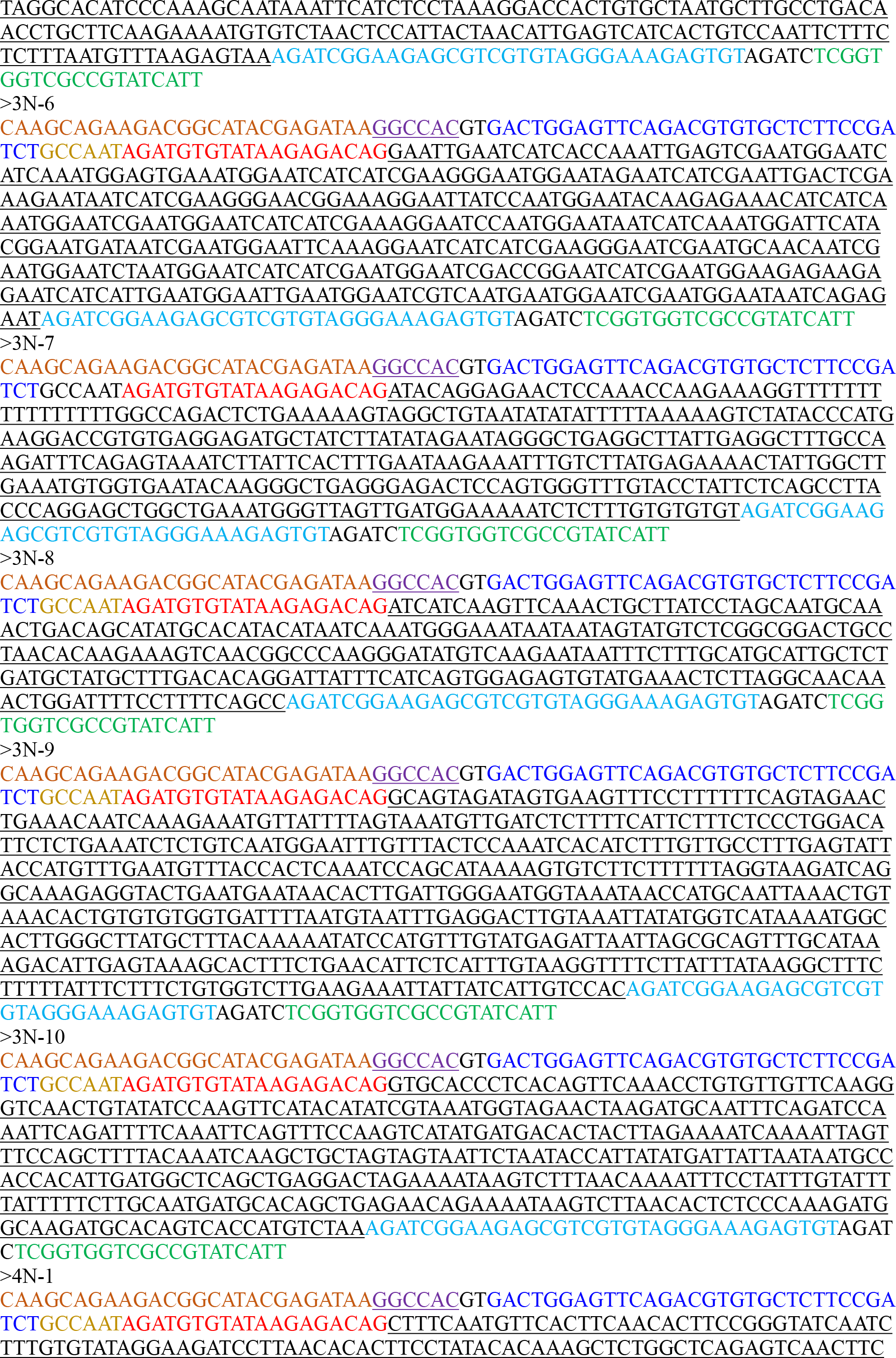

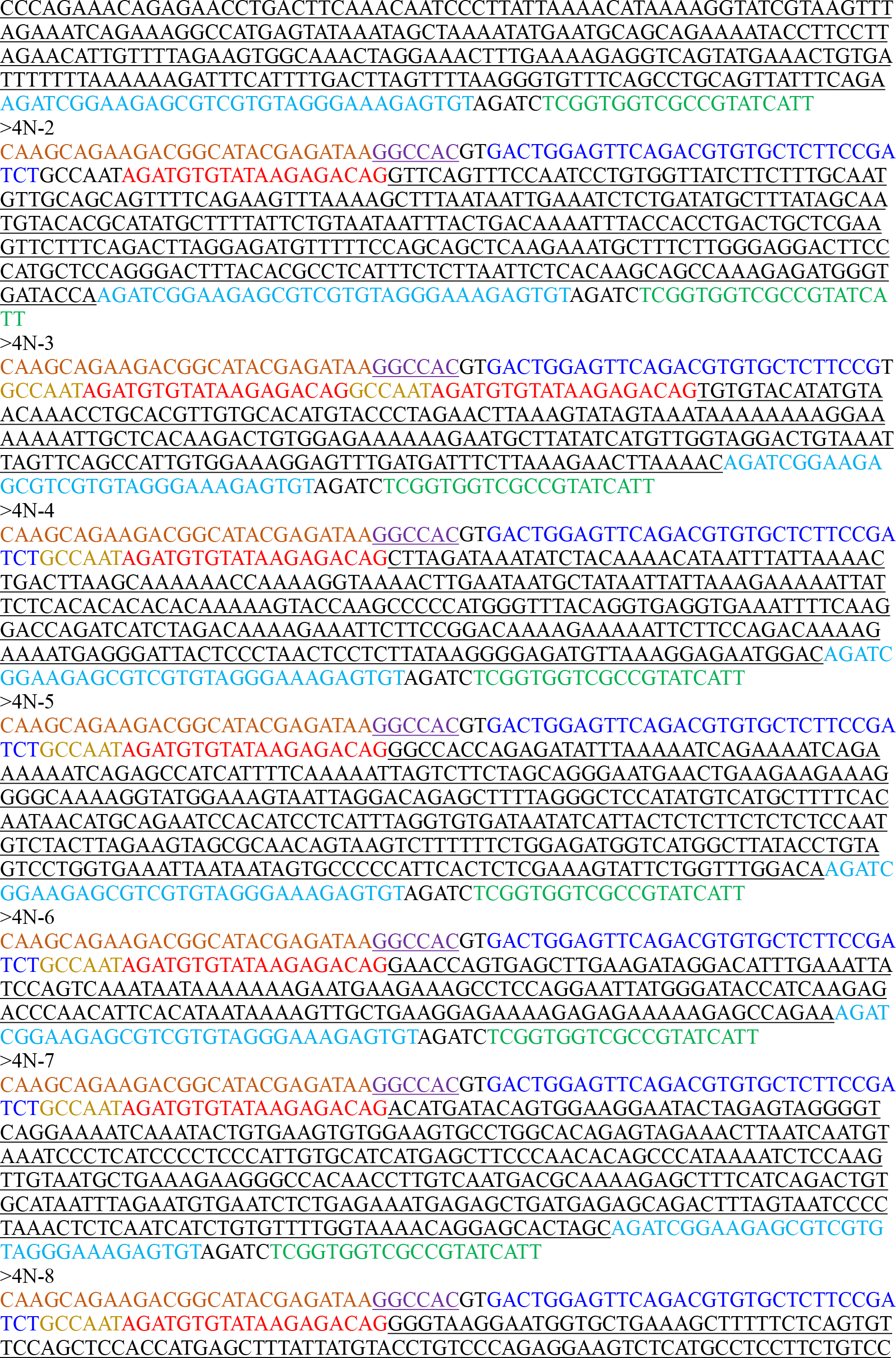

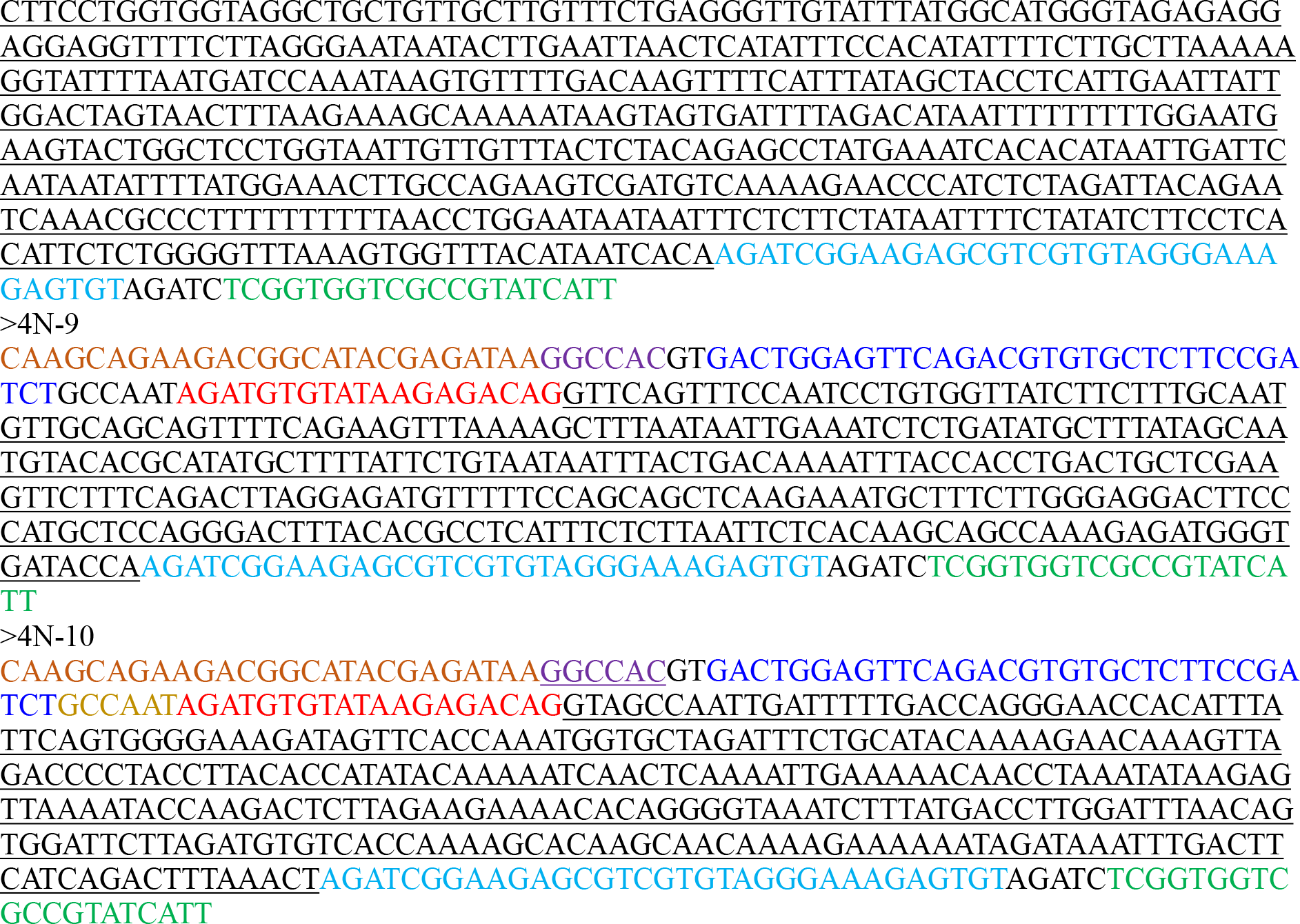

